# A Deep Neural Network Model of Audiovisual Speech Recognition Reports the McGurk Effect

**DOI:** 10.1101/2025.08.20.671347

**Authors:** Haotian Ma, Zhengjia Wang, Xiang Zhang, John F Magnotti, Michael S Beauchamp

## Abstract

In the McGurk effect, perception of an auditory syllable changes dramatically when it is paired with an incongruent visual syllable, countering our intuition that speech perception is solely an auditory process. The dominant modeling framework for the study of audiovisual speech perception is that of Bayesian causal inference, but current Bayesian models are unable to predict the wide range of percepts evoked by McGurk syllables. We explored whether a deep neural network (DNN) known as AVHuBERT could provide an alternative modeling framework. AVHuBERT model variants were presented with McGurk syllables consisting of auditory “ba” paired with visual “ga” recorded from 8 different talkers. AVHuBERT identified McGurk syllables as something other than “ba” at a rate of 59%, demonstrating a robust McGurk effect. The rate of the McGurk effect was similar to that observed in humans: 100 participants presented with the same McGurk syllables reported non-“ba” percepts on 56% of trials. AVHubert variants and humans produced a wide variety of responses to McGurk syllables, including the canonical McGurk fusion percept of “da“, responses without any initial consonant such as “ah” and responses with other initial consonants such as “fa“. The ability to predict percepts experienced by humans but not predicted by current Bayesian models suggest that DNNs and Bayesian models may provide complementary windows into the perceptual mechanisms underlying human audiovisual speech perception.

## Introduction

Humans perceive speech by integrating auditory and visual information from the face and voice of the talker. The powerful influence of visual information on speech perception is demonstrated by an illusion known as the McGurk effect, in which the pairing of incongruent auditory and visual syllables results in the perception of something other than the auditory syllable (McGurk & MacDonald, 1976). In the decades since its discovery, the McGurk effect has become a staple of psychology teaching and research, with the original paper cited nearly 10,000 times. However, mysteries remain about the perceptual processes underlying the McGurk effect.

Audiovisual speech perception, including the McGurk effect, is an example of multisensory integration, the process by which information from different sensory modalities are combined (Angelaki et al., 2009; Stein & Meredith, 1993). The dominant modeling framework for studying multisensory integration is termed *Bayesian causal inference* (Shams & Beierholm, 2022). The *Bayesian* aspect of the framework refers to the necessity of weighting the sensory cues by their reliability (Ernst & Banks, 2002), while *causal inference* refers to the necessity of estimating the likelihood that the cues arise from a single cause (Kording et al., 2007). Unlike simple audiovisual tasks, such as estimating the location of a paired beep and flash, applying Bayesian causal inference models to audiovisual speech perception is challenging because the input space (the physical characteristics of auditory and visual speech) and the output space of speech tokens do not have a simple relationship (Ma et al., 2009). This necessitates the construction of a representational space with a small number of speech tokens within which the integration computation can be performed (Aller & Noppeney, 2019; Ma et al., 2009; Magnotti & Beauchamp, 2015, 2017; Olasagasti & Giraud, 2020; Parise, 2024; Parise & Ernst, 2023).

For instance, in the McGurk effect, the pairing of visual “ga” with auditory “ba” may result in the percept of “da” (referred to as a “fusion” percept because it differs from the unisensory components) or the percept of “ba” (the auditory component of the McGurk stimulus) (Basu Mallick et al., 2015). A Bayesian causal inference model of the McGurk effect contained a representational space with only three syllables (“ba“, “da” and “ga“) organized so that “da” is intermediate between “ba” and “ga” (Magnotti & Beauchamp, 2015, 2017). Bayesian models have the advantage that every step is mathematically simple and easily interpretable. However, humans regularly perceive thousands of different words, not just a handful of syllables.

An alternative approach is provided by deep neural networks (DNNs) trained for speech recognition. Rather than restricting output to a small set of predefined syllables, DNNs can transcribe arbitrary speech into complete words. This generalizability comes at the cost of reduced interpretability, as the purpose of the computations taking place in the dozens of layers and thousands of units in DNNs is opaque. The contrasting nature of Bayesian causal inference models (high interpretability/low generalizability) and DNNs (low interpretability/high generalizability) suggests that they could be complementary tools for studying audiovisual speech perception. A key test of models is their ability to predict human perception. Within the limits of their restricted representational space, Bayesian models accurately predict human perception of McGurk syllables (Magnotti & Beauchamp, 2017). However, it is unknown whether DNNs report the McGurk effect. To answer this question, we studied a DNN known as Audiovisual Hidden-unit Bidirectional Encoder Representations from Transformers. AVHuBERT takes as input both auditory and visual speech streams and transcribes audiovisual speech with a 1.4% word error rate, similar to human performance (Shi et al., 2022).

AVHuBERT was presented with McGurk stimuli recorded from 8 different talkers, as well as a variety of other congruent, incongruent, and unisensory syllables. The responses of AVHuBERT were compared with human participants presented with the same stimuli. In addition to the fusion percept of “da” and the auditory percept of “ba“, humans report a variety of other percepts to McGurk syllables that are not predicted by current Bayesian models, including percepts of only a vowel without any initial consonant, such as “ah“, and percepts with other initial consonants, such as “fa” (Basu Mallick et al., 2015). If AVHuBERT reproduces the full range of percepts reported by humans to McGurk syllables (including the fusion percept, the auditory percept, and percepts not predicted by Bayesian models) it would indicate potential similarity with human perceptual processes, warranting further investigation of DNNs as a modeling tool to complement Bayesian causal inference models. Conversely, a failure to report human-like responses would suggest that AVHuBERT is poorly suited as a model for human perception of the McGurk effect.

## Methods

### Speech stimuli

Stimuli were recorded from 8 different talkers (4 male and 4 female) and consisted of auditory and visual recordings of the syllables “ba“, “ga” and “da“. The recordings were presented in auditory-only format (Aba; Aga; Ada); visual-only format (Vba; Vga; Vda); and audiovisual format. The audiovisual stimuli consisted of congruent pairings (AbaVba; AgaVga; AdaVda) and two incongruent pairings, the McGurk pairing (AbaVga) and the inverse pairing (AgaVba).

### Response scoring

Speech stimuli were presented to AVHuBERT using the default processing pipeline described in the AVHuBERT documentation (https://github.com/facebookresearch/av_hubert). For each stimulus, AVHuBERT generated five response predictions and the likelihood of each one.

Response likelihoods were converted to percentages so that the sum for each stimulus totaled 100%. The first phoneme of each response was classified as one of the phonemes in the Carnegie Mellon pronouncing dictionary (http://www.speech.cs.cmu.edu/cgi-bin/cmudict) as described in (Yu et al., 2024).

AVHuBERT responses to McGurk syllables contained 18 different initial phonemes. To simplify data presentation, these were grouped into 6 categories that together comprised 99% of responses to McGurk stimuli. The most frequent response was the veridical responses “B” (41% of responses), corresponding to that auditory “ba” component of McGurk syllables. The remaining responses, referred to as non-veridical, were grouped into five categories. The category of “fusion responses” consisted of the phonemes “TH” (13%), “D” (5%), “DH” (3%) and “T” (0%). The category of “responses without an initial consonant” consisted of “ah” and similar responses (classified as “AA” by the dictionary, 23% of responses), “huh” and similar responses (classified as “HH“, 3%),“uh” and similar responses (classified as “AH“, 1%), or “our” and similar responses (classified as “AW“, 3%). “F” (6%), “L” (0.8%) and “G” (0.3%) responses received their own category. The same response categories were applied for all stimuli. See the supplementary online material (SOM) for a complete list of raw responses, computed phoneme likelihoods, and analysis code.

### AVHuBERT model and model variants

The original AVHuBERT-large model (Shi et al., 2022) was downloaded from GitHub (https://github.com/facebookresearch/av_hubert, fixed version archived to read-only on Sept 1, 2024) and installed on an Apple Mac Studio with 192 GB of RAM. The AVHuBERT architecture consists of four functional parts: separate auditory and visual feature extractors; 24 layers of transformer encoders that combine the auditory and visual features; and a sequence-to-sequence decoder that produces transcribed output in the form of text tokens.

To study the sensitivity of the AVHuBERT model, slight perturbations were introduced into the first six transformer layers (the model layers most important for integrating auditory and visual speech information). The perturbations were restricted to the linear output layer (“out_proj”) of the transformer encoders (Vaswani et al., 2017). A normal distribution with mean of zero and standard deviation of 0.1 was used as the noise generator, resulting in only small changes to the model weights. One hundred unique random number seeds were used to create 100 model variants: this assured that each model variant contained different model weights, but that the variants could be uniquely identified and reproduced if necessary.

### Significance calculations

To calculate statistical significance, t-tests were performed using *R*. See the supplementary online material (SOM) for full statistical results.

### Comparison with human responses

For comparison with the AVHuBERT responses to McGurk stimuli, we reanalyzed data from 100 human participants collected for the open choice portion of Experiment 2 of Basu Mallick et al. (2015). Identical McGurk stimuli were tested in humans and in AVHuBERT model variants, and the same response scoring procedure (described above) was applied. Human responses to the incongruent audiovisual syllable AgaVba are from (Magnotti & Beauchamp, 2017).

## Results

In the McGurk effect, pairing auditory “ba” with an incongruent visual “ga” results in a non-veridical percept (something other than “ba“). For AVHuBERT model variants presented with McGurk syllables recorded from 8 different talkers, non-veridical responses were observed at a rate of 59% (Figure 1; means across recordings and model variants). Human participants presented with the same stimuli reported non-veridical percepts at a similar rate (56%; *p* = 0.3 from unpaired *t*-test with unequal variance).

**Figure 1.**
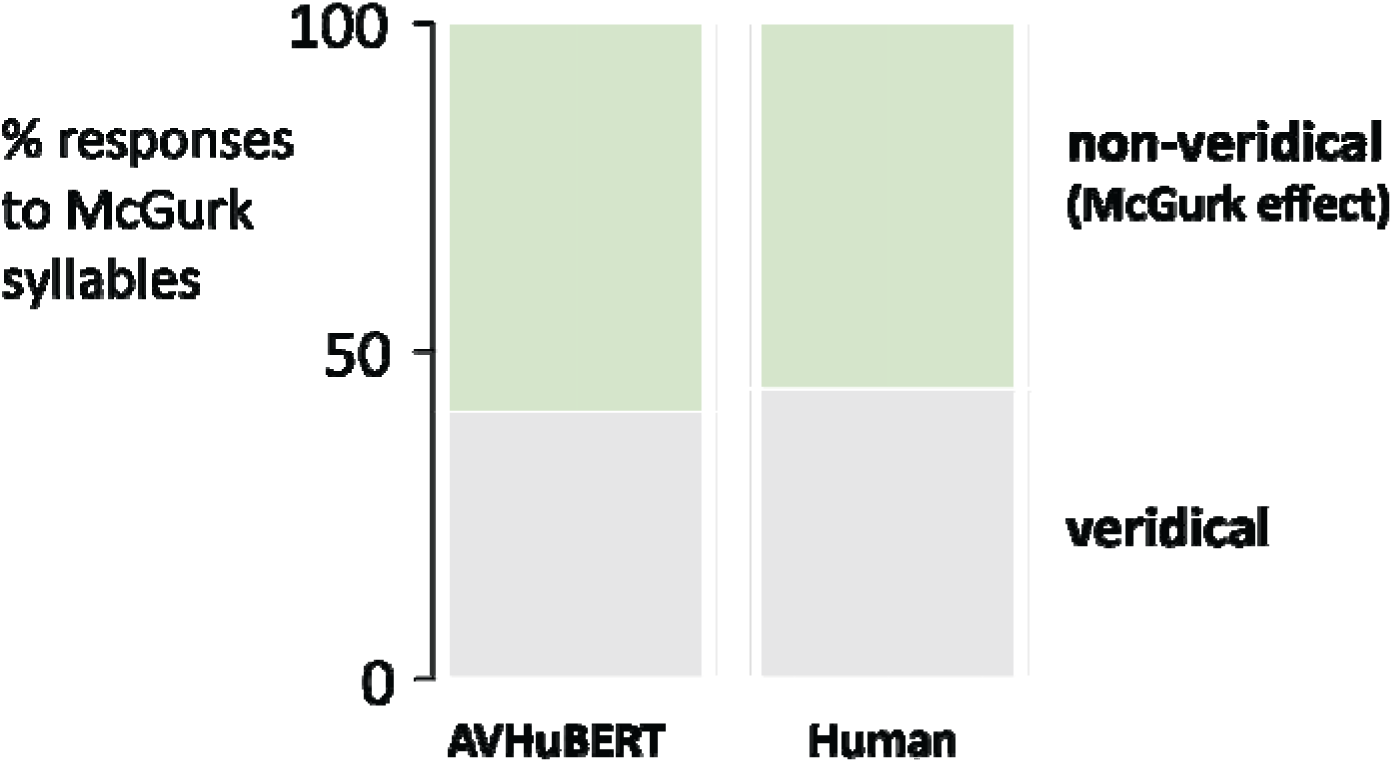
McGurk syllables (auditory “ba” paired with visual “ga“) recorded from 8 different talkers were presented to 100 AVHuBERT model variants and 100 human participants. Veridical perception was defined as a response of “B“, the auditory component of the syllable (gray shaded region). Non-veridical perception, indicating the McGurk effect, was defined as all responses other than “B” (green shaded area). Mean values across recordings and variants/participants.

AVHuBERT’s responses to McGurk syllables were examined in more detail (Figure 2A). The most-frequent non-veridical responses contained *no initial consonant* (such as “AA“), accounting for 50% of non-veridical responses (30% of all responses). The second most-frequent were *fusion responses* (such as “TH” and “D“), accounting for 36% of non-veridical responses (21% of all responses). The third most-frequent were responses with the initial consonant “F” (9% of non-veridical responses, 6% of all responses); all other responses occurred with less than 1% frequency.

**Figure 2.**
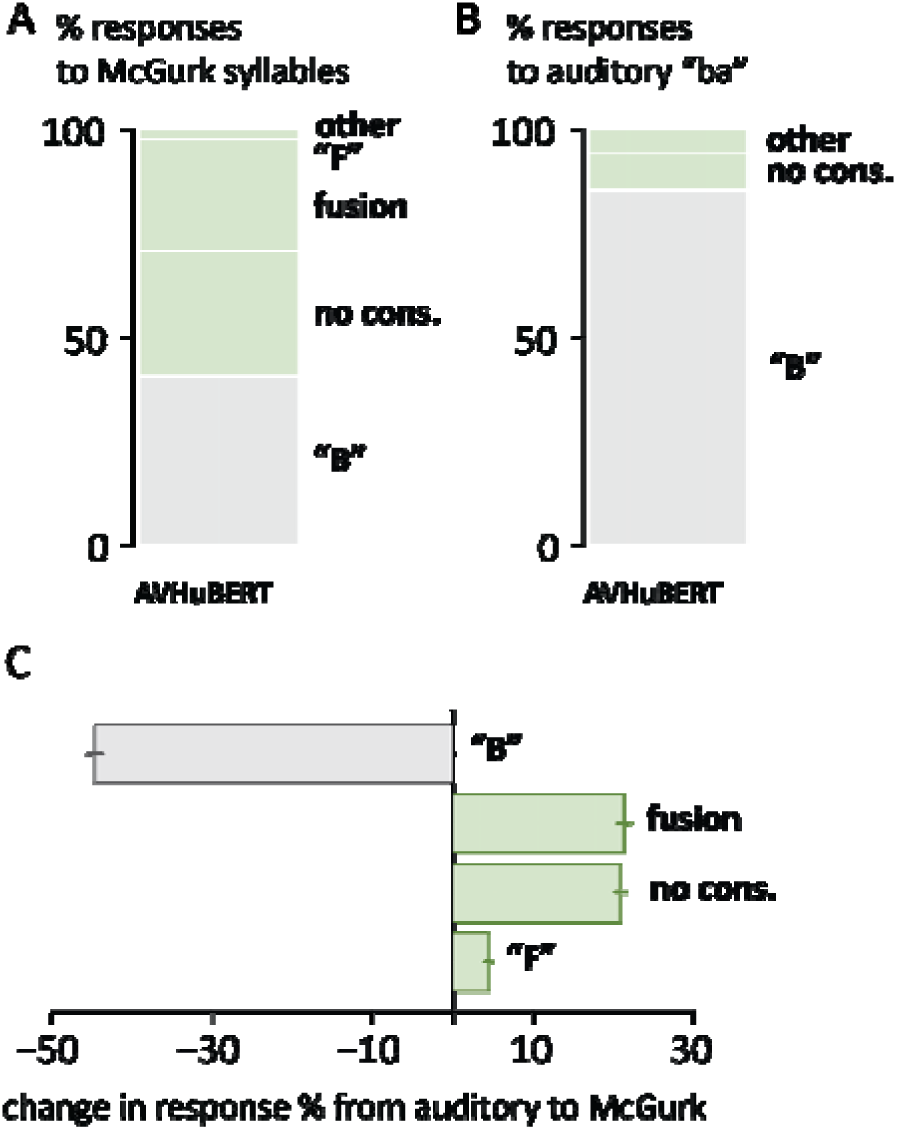
**A.** McGurk syllables (auditory “ba” paired with visual “ga“) were presented to 100 AVHuBERT model variants. Non-veridical perception, indicating the McGurk effect, was defined as all responses other than “B” (green shaded areas). “No-cons.” includes all responses with no initial consonant, such as “AA“. “Fusion responses” include “D” and related phonemes. Mean values across recordings and variants. **B.** Responses of AVHuBERT model variants to auditory “ba” presented on its own. **C.** Change in response rates between auditory “ba” and McGurk syllables. The rate of “B” responses decreased with a corresponding increase in the rate of other response categories. Bars for “G” (+0.3%) and “L” (+0.8%) responses not shown. Mean across recordings and variants, SEM across variants.

AVHuBERT responses to McGurk syllables were very different than responses to auditory “ba” presented on its own, demonstrating a strong influence of the incongruent visual “ga” in the McGurk syllables (Figure 2B, 2C). The largest change was in the rate of veridical responses (initial phoneme “B“), which decreased by 45% in the presence of incongruent visual “ga“, from 85.2% for auditory-only “ba” to 40.7% for McGurk syllables (*p* < 10^-16^ from paired *t*-test). This was balanced by increases in *fusion responses* (from 0% to 21%, *p* < 10^-16^) and responses with *no initial consonant* (from 9% to 30%, *p* < 10^-16^). Response increases were also observed for “F” (1% to 6%, *p* < 10^-16^), “L” (0% to 0.8%, *p* = 10^-5^) and “G” (0% to 0.3%, *p* = 10^-4^).

*Fusion responses* were observed at significantly higher rates for McGurk syllables (21%) than any of the control stimuli, including the congruent pairings of auditory “ba” with visual “ba” (AbaVba; 0%) and AgaVga (0.3%); the incongruent pairing of AgaVba (0.1%); or the unisensory components of McGurk syllables Vga (6%) and Aba (0.2%); all *p* < 10^-16^. In contrast, responses with *no initial consonant* were observed at rates of 58% for Vga and 9% for Aba, indicating that these responses were not specific to McGurk syllables, which evoked them at a rate of 30%.

### Comparisons with human perception

The mean rate of non-veridical responses to McGurk syllables was similar for AVHuBERT model variants and human participants, 59% *vs.* 56%. As shown in Figure 3A, there was high variability in the rate of non-veridical responses across different AVHuBERT variants (range 38% to 85%) and human participants (0% to 100%). A wide variety of different initial phonemes were observed in the responses to McGurk syllables: AVHuBERT responses contained 18 different initial phonemes, while human responses contained 17 different initial phonemes (Figure 3B). Humans had more fusion responses than AVHuBERT (41% *vs.* 21%, *p* = 10^-8^) and more “G” responses (6% *vs.* 0.3%, *p* = 10^-6^) while AVHuBERT had more responses with no initial consonant (30% *vs.* 3%, *p* < 10^-16^) and more “F” responses (6% *vs.* 2%; *p* = 10^-4^). Response rates for “L” were similar (1.3% human *vs.* 0.8% AVHuBERT, *p* = 0.3).

**Figure 3.**
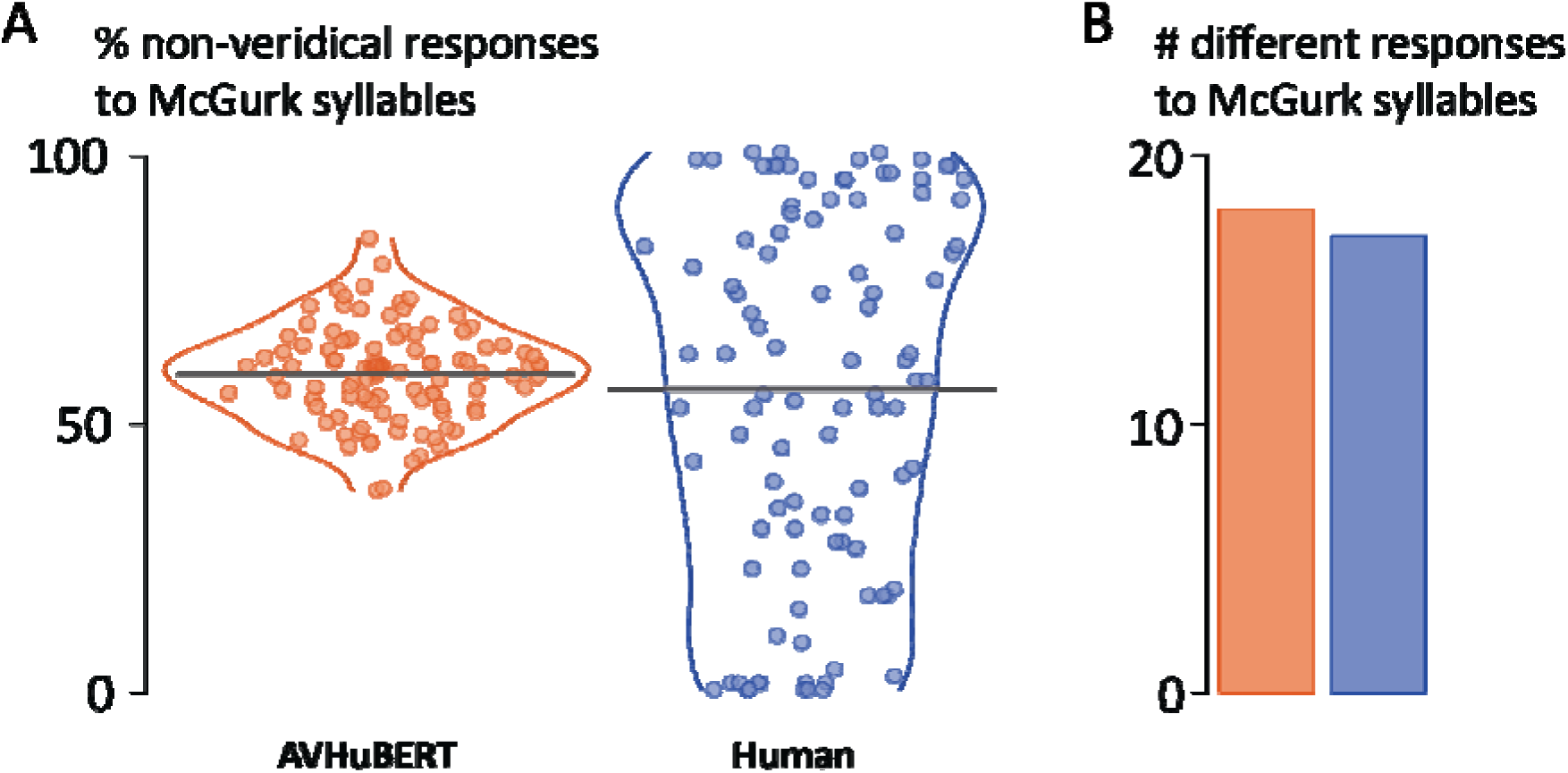
**A.** The rate of the McGurk effect, defined as non-veridical (other than “B“) responses to McGurk syllables. Orange symbols show results for 100 AVHuBERT model variants (larger symbol near center of plot shows value for original AVHuBERT). Blue symbols show results for 100 human participants. One symbol per variant/participant, mean across 8 recordings for each variant/participant. Black line shows mean across variants/participants. Horizontal spread of symbols and colored outline indicates probability density. Shaded area shows one SD from the mean. **B.** The total number of different initial phonemes across all responses to McGurk syllables for all AVHuBERT model variants (orange) and human participants (blue).

In contrast with the variety of responses observed for McGurk syllables, responses were relatively uniform for the congruent syllables AbaVba, AgaVga and AdaVda, with human participants and AVHuBERT variants producing correct responses at a rate of 97% for humans and 94% for AVHuBERT (means across syllables, recordings and participants/variants). The inverse pairing of auditory “ga” paired with incongruent visual “ba” produced mainly “G” responses in both humans (96%) and AVHuBERT variants (91%), demonstrating that the variety of responses observed for McGurk syllables was not solely attributable to audiovisual incongruence.

## Discussion

The signature of the McGurk effect is a change in the perception of auditory “ba” when paired with an incongruent visual “ga“. When presented with McGurk syllables, AVHuBERT generated a non-veridical report on 59% of trials, providing an affirmative answer to the question “Does AVHuBERT report the McGurk effect?” Humans presented with McGurk syllables reported a similar rate of non-veridical percepts. Both AVHuBERT and humans produced a wide variety of responses to McGurk syllables, including the fusion percept of “da” and related phonemes described in the original description of the effect (McGurk & MacDonald, 1976).

Fusion responses were specific to McGurk syllables, occurring at significantly higher rates for McGurk syllables than for any of the tested control stimuli, including the auditory and visual components of McGurk syllables presented on their own or the inverse pairing of visual “ba” and auditory “ga“.

The ability to predict human behavior is an important requirement for models, and the similarities between human responses and AVHuBERT suggest that it could be a useful tool for interrogating the perceptual processes underlying human audiovisual speech perception.

Most current models of the McGurk effect operate in the framework of Bayesian causal inference (Aller & Noppeney, 2019; Magnotti & Beauchamp, 2015, 2017; Olasagasti & Giraud, 2020; Parise, 2024; Parise & Ernst, 2023) making it worthwhile to compare Bayesian and DNN models.

Bayesian causal inference models require explicit assumptions about the representational space used by humans during speech perception. The original description of the McGurk effect proposed an explanation grounded in perceptual processing: “da” is pronounced with an open mouth, compatible with the visual properties of “ga”, while in the auditory domain, the acoustics of “da” are similar to “ba”. Thus, the fusion percept represents a compromise between conflicting sensory cues (McGurk & MacDonald, 1976). The CIMS (causal inference of multisensory speech) Bayesian model instantiated this idea by assuming a two-dimensional representational space with auditory features on one axis and visual features on the other axis. In the representational space, “da” is intermediate between “ba” and “ga”. CIMS accurately models human perception of McGurk syllables and the inverse syllable AgaVba (Magnotti & Beauchamp, 2017).

One way to conceptualize differences between Bayesian causal inference and DNNs are along the axes of generalizability and interpretability. Bayesian causal inference models generalize poorly. They can only return the tokens in the representational space with which they are constructed, such as the three syllables in the two-dimensional representational space of the CIMS model (Magnotti & Beauchamp, 2017). However, modeling studies of words in noise suggest that audiovisual speech is very high-dimensional with 20, 50 or more dimensions (Ma et al., 2009). No one has yet been able to create a Bayesian model that parametrizes these dimensions to integrate arbitrary auditory and visual speech. Therefore, current Bayesian models can only predict human perception in the context of a forced choice experimental design in which the forced choices correspond to the modeled syllables. Of course, human speech perception is not limited to a handful of syllables and the inability to generalize to arbitrary audiovisual speech limits the utility of Bayesian models.

Unlike Bayesian models, DNNs such as AVHuBERT generalize well. By design, they can transcribe any auditory, visual or audiovisual speech and estimate the likelihood of the component words. The generalizability comes at the cost of interpretability. AVHuBERT has 325 million parameters to describe the dozens of layers in its DNN, making the computations performed within it difficult to discern (Shi et al., 2022). Conversely, Bayesian causal inference models are parsimonious, with a few steps controlled by a handful of parameters. Each computation is mathematically simple and can be understood conceptually as the weighted average of alternative percepts.

AVHuBERT frequently classified the visual component of McGurk syllables (Vga) as “no-initial-consonant” and there were many “no-initial-consonant” responses to McGurk syllables. These “no-initial-consonant” responses may therefore represent a visual-corresponding response for AVHuBERT. Conversely, the rates of “G” responses (corresponding to the true visual component) and fusion responses were significantly lower in AVHuBERT than humans. The combination of higher rates of some non-veridical responses and lower rates of others resulted in similar overall rates of non-veridical responses for AVHuBERT and humans.

One recent discovery about the McGurk effect is the existence of high individual variability, a finding that has been replicated across speakers of different native languages (Basu Mallick et al., 2015; Dong et al., 2025; Magnotti et al., 2024, 2025; Strand et al., 2014; Tiippana et al., 2023). As shown in Figure 2B, human participants span the entire range of visual influence, from 0% to 100%, despite having normal or corrected-to-normal vision.

In the present study, AVHuBERT model variants were constructed by adding small amounts of noise to the model connections in the DNN layers most important for audiovisual integration. This produced a four-fold variation in visual influence for McGurk syllables (15% to 62%) while retaining high accuracy for congruent audiovisual syllable.

The variation across AVHuBERT variants in the rate of the McGurk effect was less than the variation across humans. To increase AVHuBERT variation, one approach would be to retrain or fine-tune AVHuBERT with different speech material. Humans experience very different language environments during development and simulating this process by training DNN models with speech that varied in the number of audible and visible talkers, the number of talkers speaking at once, and the levels of background noise, would be expected to increase the variability across models.

The architecture of DNNs was inspired by biological neural systems, and therefore there is a possible correspondence between them (*e.g.* model units ∼ neurons, model weights ∼ synaptic strengths). Examining the internal structure of AVHuBERT might give clues to the neural substrates of the McGurk effect, as has been successfully demonstrated for DNNs and convolutional neural networks (CNNs) trained to accomplish visual, auditory, and language tasks (Kell et al., 2018; Kumar et al., 2024; Li et al., 2023; Saddler & McDermott, 2024; Yamins et al., 2014; Yamins & DiCarlo, 2016). Of course, the many possible methodological confounds in comparisons between DNN/CNNs and human brains warrant careful consideration (Dujmovic et al., 2024; Hadidi et al., 2025) and emphasize the diversity of possible neural network architectures (Rideaux et al., 2021). In the perceptual domain, the ability of AVHuBERT to transcribe any speech stimulus means that it should be possible to test combinations of incongruent audiovisual speech that have never been presented to humans before, potentially leading to the discovery of previously unknown illusions related to the McGurk effect.

## Declarations

### Ethics Approval

The study was approved by the University of Pennsylvania IRB.

### Consent to participate

Informed consent was obtained from all participants included in the study.

### Consent for publication

Participants gave written informed consent for publishing the overall results prior to the experiment.

### Availability of data and materials

All data reported in this manuscript are included as supplemental material.

### Code availability

All analysis code is included as supplemental material.

### Open Practices Statement

All data and analysis code necessary to reproduce the results in this manuscript are included as supplemental material.

